# Kinome rewiring reveals AURKA is a molecular barrier to the efficacy of PI3K/mTOR-pathway inhibitors in breast cancer

**DOI:** 10.1101/158295

**Authors:** Hayley J Donnella, James T Webber, Rebecca S Levin, Roman Camarda, Olga Momcilovic, Nora Bayani, James Korkola, Kevan M Shokat, Andrei Goga, John Gordan, Sourav Bandyopadhyay

**Affiliations:** University of California, San Francisco. Department of Bioengineering and Therapeutic Sciences, San Francisco, CA 94148.; University of California, San Francisco. Department of Cellular and Molecular Pharmacology, San Francisco, CA 94148.; University of California, San Francisco. Department of Medicine, San Francisco, CA 94148.; Oregon Health and Sciences University. Center for Spatial Systems Biology, Portland, OR 97239; Howard Hughes Medical Institute.

## Abstract

Dysregulation of the PI3K-AKT-mTOR signaling network is a prominent feature of breast cancers. However, clinical responses to drugs targeting this pathway have been modest. We hypothesized that dynamic changes in signaling, including adaptation and feedback, limit drug efficacy. Using a quantitative chemoproteomics approach we mapped dynamic changes in the kinome in response to various agents and identified signaling changes that correlate with drug sensitivity. Measurement of dynamics across a panel of breast cancer cell lines identified that maintenance of CDK4 and AURKA activity was associated with drug resistance. We tested whether incomplete inhibition of CDK4 or AURKA was a source of therapy failure and found that inhibition of either was sufficient to sensitize most breast cancer cells to PI3K, AKT, and mTOR inhibitors. In particular, drug combinations including the AURKA inhibitor MLN8237 were highly synergistic and induced apoptosis through enhanced suppression of mTOR signaling to S6 and 4E-BP1 leading to tumor regression *in vivo.* This signaling map identifies survival factors whose presence limits the efficacy of target therapy and indicates that Aurora kinase co-inhibition could unlock the full potential of PI3K-AKT-mTOR pathway inhibitors in breast cancer.

## INTRODUCTION

Mutations and aberrant signaling of the PI3K–AKT–mTOR pathway (PI3K-pathway) is a prominent feature of breast and many other cancers. Genomic alterations of PI3K-pathway components, including *PTEN, PIK3CA,* and *AKT1* occur in over 60% of breast malignancies^1^. Despite these data, multiple clinical trials demonstrate that monotherapy responses to agents directed against this pathway are modest^2–5^. The clinical observation that breast cancers fail to respond to therapy suggests that additional factors modulate cellular response and drive resistance. A prominent feature of this pathway is drug-induced signaling adaptations and feedback mechanisms resulting in suboptimal drug responses^2, 6–8^. Therefore, it is likely that understanding and targeting these dynamic changes in signaling will be important in optimizing this class of agents. Emerging evidence indicates that adaptive changes in response to a wide variety of targeted therapies may also be critical for mediating tumor cell survival during treatment^9–12^, thus highlighting the need to understand the dynamics of response at the level of signaling networks in order to identify more effective treatments.

In principle, the measurement of dynamic changes elicited by therapy can be used to develop novel drug combinations. Such efforts have largely focused on adaptive signaling changes leading to pathway reactivation, thereby causing drug resistance^9, 13–16^. An alternative approach could be to identify survival factors whose presence limits cellular dependence on the targeted pathway. Conceptually, the maintenance of such factors during drug treatment could provide survival signals facilitating drug resistance and since failure to inhibit survival factors would limit the efficacy of targeted therapy, their identification could reveal new synthetic lethal strategies to enhance therapeutic responses. Previous identification of such factors have led to the understanding that the activation of apoptotic machinery^14, 17, 18^ and impairment of protein synthesis^15^ is required for sensitivity to a wide variety of drugs. In the context of breast cancer, multiple efforts in the field have identified mTORC1 as a survival factor whose suppression is necessary for PI3K-pathway inhibitor sensitivity^17, 19, 20^. This has led to clinical trials combining PI3K and mTOR inhibitors, yet reported clinical results have yielded suboptimal outcomes due to increased systemic toxicity and cytostatic tumor effects^21^. Hence, there remains a pressing need to uncover new combination targets in order to improve therapeutic efficiency of PI3K-pathway inhibitors. Identifying additional survival factors will require a comprehensive understanding of signaling dynamics in response to treatment and an understanding of how these dynamics contribute to drug resistance.

Little is known about global kinome rewiring in response to targeted therapies and the impact this has on drug resistance. This is due in part to the limited technologies available for capturing kinome activity, which are restricted by available antibody reagents and the narrow utility of phosphoproteomics studies to infer kinase activity from substrates. Recently, a kinase enrichment strategy has been developed using a chemoproteomics technique that combines kinase affinity capture with quantitative mass spectrometry (MS). This approach uses a multiplexed set of type I kinase inhibitors immobilized onto beads (MIBs), which are used to affinity purify a diverse set of active kinases through their increased avidity for ATP compared to inactive kinases. Enriched kinases are then identified and quantified by LC MS/MS (MIBs/MS), enabling simultaneous measurement of many endogenous kinases based on their activity state and abundance^22–25^. In contrast to previous studies^9, 26, 27^, increasing the scale of this approach could be used to generate a quantitative map of kinase dynamics thus allowing for global comparisons of cellular responses to a variety of agents.

Here we applied the MIBs/MS approach to identify signaling changes associated with drug efficacy by mapping the kinome following exposure to targeted therapies across a panel of breast cancer cell lines of various subtypes and genotypes. To illustrate the utility of this signaling map, we compared kinome activity profiles between drug-sensitive and resistant cells to generate a kinome-response signature associated with drug sensitivity. By performing a systematic analysis of signaling dynamics following drug treatment, we identified that failure to inhibit a set of kinases including CDK4 and AURKA was associated with drug resistance to a diverse set of targeted therapies. Further analysis revealed that inhibition of CDK4 or AURKA was sufficient to engender strong synergistic responses when combined with inhibitors of PI3K, AKT, and mTOR. This provides an effective new framework for the unbiased identification of survival factors acting as molecular barriers to the efficacy of drugs, and we demonstrate the utility of this approach by developing rational combination strategies to enhance responses to PI3K-pathway inhibitors in breast cancer.

## RESULTS

### Generation of a kinome map and identification of dynamic changes that correlate with drug sensitivity

We applied an unbiased proteomic strategy to measure kinome rewiring in response to drug treatment. Kinome profiling was performed via a chemoproteomics approach using multiplexed inhibitor beads (MIBs) coupled with mass spectrometry (MIBs/MS)^22–24^. Multiplex inhibitor beads (MIBs) consist of a mixture of sepharose beads covalently linked to 12 kinase inhibitors ranging from moderately selective (e.g. Lapatinib, Sorafenib) to pan-kinase inhibitors (e.g. Purvalanol B, Staurosporine) for broad kinome coverage (Fig. 1a and Supplementary Fig. 1). Because type I kinase inhibitors preferentially bind kinases in their active conformation, kinase capture by MIBs is a function of kinase expression, the affinity of kinases for the immobilized inhibitors, and the activation state of the kinase^28^. In this approach, vehicle or drug treated cell lysates were incubated with MIBs, and enriched kinases were eluted, digested into peptides, and subjected to identification and quantification by LC MS/MS using a label-free quantitation pipeline (see Methods)^22^. The ability to detect a kinase within a given cell line is a function of the inhibitor binding spectrum, modified by the level of expression, and we estimate that the current approach is able to capture roughly 35% of highly expressed kinases in a given sample (Supplementary Fig. 2).

**Figure 1.**
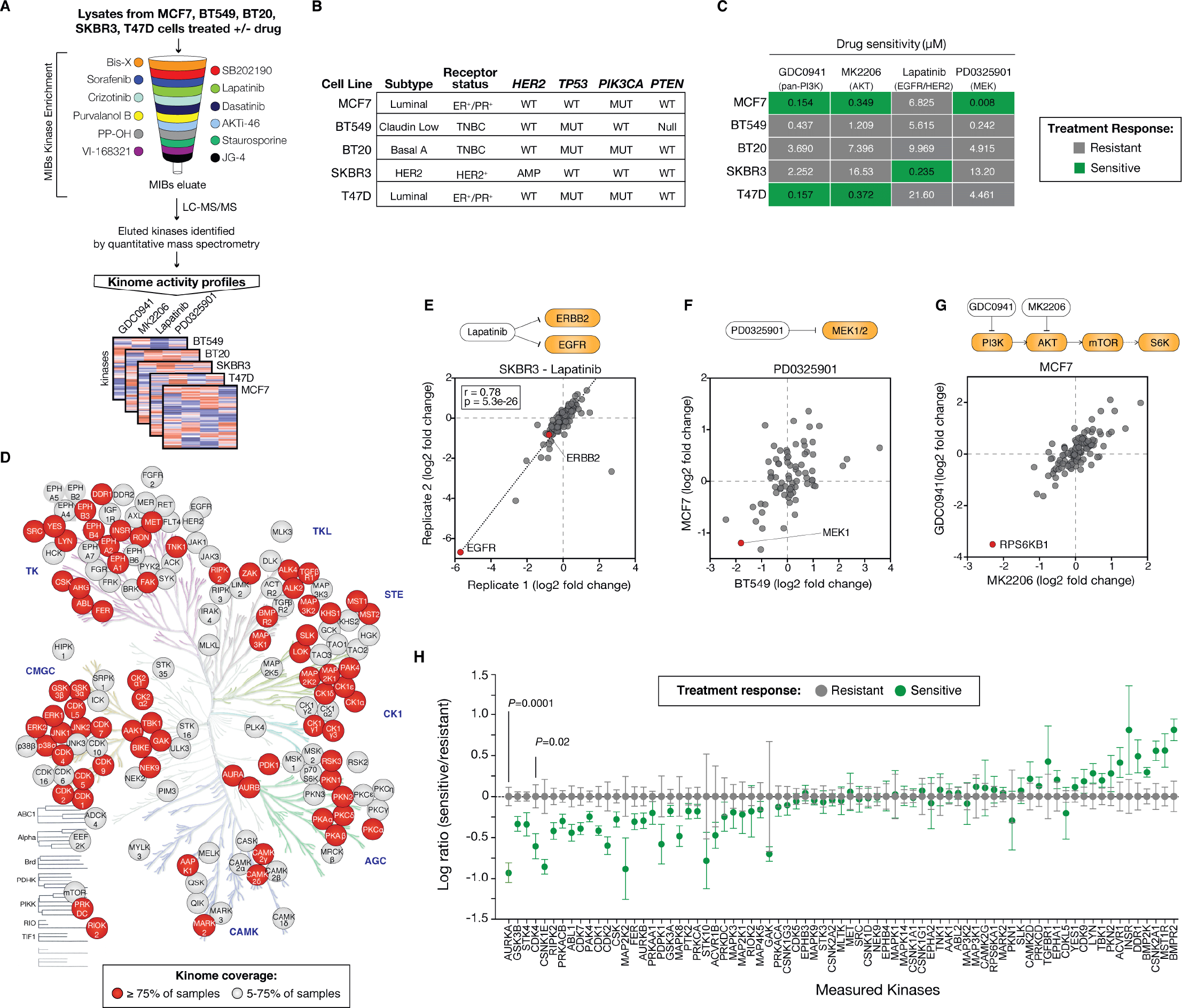
Measurement of kinome dynamics to identify correlates of drug sensitivity. (**a**) Schematic of approach using multiplex inhibitor beads followed by mass spectrometry (MIBs/MS). Sample lysates are passed through a column containing the indicated kinase inhibitors covalently linked to beads. After washing, bound proteins are eluted, trypsin digested and quantified through label-free mass spectrometry. (**b**) Table of breast cancer cell lines used in this study with the indicated molecular subtypes and mutational status. (**c**) Drug sensitivity of cell lines indicated by IC_50_ and categorized into relative drug sensitivity or resistance. Because IC_50_ was not reached for PD0325901, IC_25_ is shown. (**d**) Human kinome tree annotated with kinases identified in this study and colored based on the percentage of total samples where each particular kinase could be quantified. (**e**) Comparison of kinase activity scores, expressed as a ratio of measured kinases in SKBR3 cells treated with 200nM Lapatinib versus DMSO treated control across biological replicates. Pearson correlation and p-value indicated. (**f**) Comparison of kinase activity ratios in BT549 and MCF7 cells treated with 100nM PD0325901. For clarity, one outlier kinase (GAK, BT549 log_2_ fold change 8.3) was removed. (**g**) Comparison of kinase activity ratios for MCF7 cells treated with either 250nM MK2206 or GDC-0941. (**h**) Categorical analysis of kinome dynamics that occur in drug sensitive treatment responses (n = 6) versus resistant treatment responses (n = 14) for all drugs pooled together. For visualization purposes, each kinase was centered on the mean of resistant samples. Data shown for 75 kinases which could be measured in >75% of samples. All drug treatments are for 24 hours. Error bars are s.e.m. *P* values calculated using a two-sided t-test.

We applied this strategy to a panel of breast cancer cell lines of various subtype and genotype classifications, and measured kinome dynamics following treatment with an array of targeted therapies. Cell lines were chosen to maximize diversity and span the major subtypes of breast cancer, and display large differences in transcriptional profiles (Supplementary Fig. 3). All lines harbored mutations in PI3K-pathway genes including *PIK3CA-mutant* MCF7 (ER^+^/PR^+^), BT20 (receptor negative) and T47D (ER^+^/PR^+^); PTEN-null BT549 (receptor negative); and HER2-amplified SKBR3 (HER2^+^) (Fig. 1b). Cell lines were treated for 24 hours with DMSO or kinase inhibitors relevant to breast cancer signaling including the EGFR/HER2 inhibitor Lapatinib (200nM), the pan-Class I PI3K inhibitor GDC-0941 (250nM), the AKT inhibitor MK2206 (250nM), and the MEK inhibitor PD0325901 (100nM) (Fig. 1c). All together, MIBs/MS was able to quantify changes in 151 kinases, and robustly measured 75 kinases which were present over 75% (15/20) of samples (Fig. 1d, Supplementary Table 1). Significant drug-induced changes were detected in 99 kinases at *p* < 0.001 corresponding to 66% of kinases measured, indicating that the drugs had widespread and significant impacts on global kinome dynamics.

To assess the quality and reproducibility of the MIBs/MS data, we initially compared biological replicates of SKBR3 (HER2^+^) cells treated with the dual EGFR/HER2 small-molecule inhibitor Lapatinib. We observed a high correlation of 0.78 between scores (defined as the log_2_ fold change compared to DMSO treatment) for identified kinases (*p* = 5e-26) (Fig. 1e). The MIBs/MS screening strategy also accurately captured activity inhibition of direct drug targets indicated by the significant decrease in scores, defined as log2 fold change compared to DMSO, for EGFR (−5.8, *p* = 6e-5) and HER2 (−0.7, *p* = 1e-4) after treatment with Lapatinib (Fig. 1e). We observed a decrease in MEK1 activity upon treatment with the MEK inhibitor PD0325901 in BT549 and MCF7 cells (log_2_ fold change score = −1.8 and −1.2, respectively, Fig. 1f). We also observed indirect pathway-specific events such as a decrease in the activity of the mTOR effector kinase RPS6KB1 when treated with either the PI3K inhibitor GDC-0941, or AKT inhibitor MK2206 in MCF7 cells (−3.5 and −2.3, respectively) (Fig. 1g). These results highlight the reproducibility of the MIBs/MS approach as well as its ability to identify direct and indirect drug targets based on reductions in both activity and abundance.

We hypothesized that the identification of shared responses across lines and drugs may lead to a more robust understanding of signaling dynamics, as opposed to changes specific to a particular drug or cell type. We therefore sought to identify dynamic changes in cells that were associated with treatment sensitivity or resistance in response to the drugs in our panel. For this purpose, we compared dose response curves and relative sensitivities across the screening set and classified each cell line as sensitive or resistant to the drugs in our panel. (Fig. 1c, Supplementary Fig. 4a-d). Normalized scores for each kinase were compared between these sensitive and resistant classifications for all drugs pooled together to identify candidate kinases whose inhibition was associated with drug sensitivity (Fig. 1h). This analysis revealed that suppression of 12 kinases was significantly associated with drug sensitivity independent of the agent used (*p* < 0.05). Among the identified candidates were kinases involved in cell cycle processes including mitotic kinases AURKA (*p* = 0.002) and CDK1 (*p* = 0.04), and kinases involved in interphase, CDK4 (*p* = 0.02) and CDK2 (*p* = 0.05). Other identified candidates included kinases associated with WNT signaling (GSK3B, *p* = 0.005 and CSNK1E, *p* = 0.02) and YAP signaling (STK4, *p* = 0.01). Beyond the diversity of pathways identified, two additional lines of evidence led us to believe that these results were specific and not linked to general impairment of the cell cycle *per se.* First, the association with sensitivity was specific for these proteins and was not linked to other cell cycle kinases. For example, we observed no correlation with sensitivity for many other cyclin-dependent kinases (CDKs) measured in our screen such as CDK6, a closely related CDK to CDK4. Second, while AURKA was significantly associated with sensitivity, its closely related paralog AURKB was not, even though it is regulated during mitosis in a similar manner^29^ (Fig. 1h). Since this approach is sensitive to the classification of sensitive and resistant treatment responses, we also performed a similar analysis based on a three-response categorization (i.e., sensitive, moderately sensitive, and resistant) and found that these results were largely robust with respect to a more granular analysis (Supplementary Fig. 4e-h). Based on the specificity we observed and the availability of clinically relevant targeted and specific inhibitors, we pursued CDK4 and AURKA for further studies. In both cases, we confirmed that maintenance of activity was associated with resistance regardless of the inhibitor used indicating that a single drug was not driving the observed trend (Supplementary Fig. 4i,j). To summarize, by performing a systematic screen of signaling dynamics following drug exposure, we identified a set of specific kinases whose maintenance was associated with resistance to drug treatment in breast cancer.

### Maintenance of CDK4 and AURKA is associated with resistance to PI3K and AKT inhibition

Given the central importance of the PI3K pathway in breast cancer, we focused our validation of molecular correlates of drug sensitivity on this pathway and expanded our analysis to related agents. Analysis of dynamic responses across the MIBs screening panel revealed a significant association between maintenance of CDK4 and AURKA activity and drug resistance (Fig. 2a,b). To confirm this result, we measured the molecular responses to treatment with the pan-PI3K inhibitor GDC-0941 in two sensitive (T47D and MCF7, IC_50_ < 200nM) and two resistant (HCC38 and MDAMB453, IC_50_ > 40*μ*M) breast cancer cell lines. A critical output of the PI3K-pathway is the activation of the mTORC1 complex, whose inhibition is necessary for sensitivity to PI3K inhibitors^19^. Consistent with this, over a 24-hour time course we observed a suppression of mTORC1 activity only in sensitive cells, as evidenced by a decrease in phosphorylated S6 at serine 240/244 downstream of mTORC1. CDK4 is a cell cycle kinase that regulates G1/S transition through the phosphorylation of RB and subsequent release of E2F target genes responsible for progression through G1^30^. Consistent with our proteomic data, we observed a modest decrease in the abundance of total CDK4 protein levels in sensitive cells, whereas resistant cells maintained their CDK4 levels throughout the course of treatment (Fig. 2c, Supplementary Fig. 5a). We next examined CDK4 activity via immunoblotting for its substrate RB and found that phospho-RB levels decreased in sensitive cells only, indicating a loss of CDK4 activity despite largely stable levels, consistent with the activity dependence of the MIBs/MS data (Supplementary Fig. 5b). We also sought to determine if the loss of CDK4 activity in drug sensitive cells was specific for PI3K inhibitors, or could be generalized to AKT inhibitors representing the next step in the PI3K-pathway. Using the AKT inhibitor MK2206, we treated resistant and sensitive lines and observed a decrease in both CDK4 abundance and phosphorylated RB in sensitive cells, while resistant cells showed no change in CDK4 or phospho-RB (Supplementary Fig. 5c-e). Therefore, these results confirm that suppression of CDK4 activity is correlated with sensitivity to the PI3K inhibitor GDC-0941 and the AKT inhibitor MK2206.

**Figure 2.**
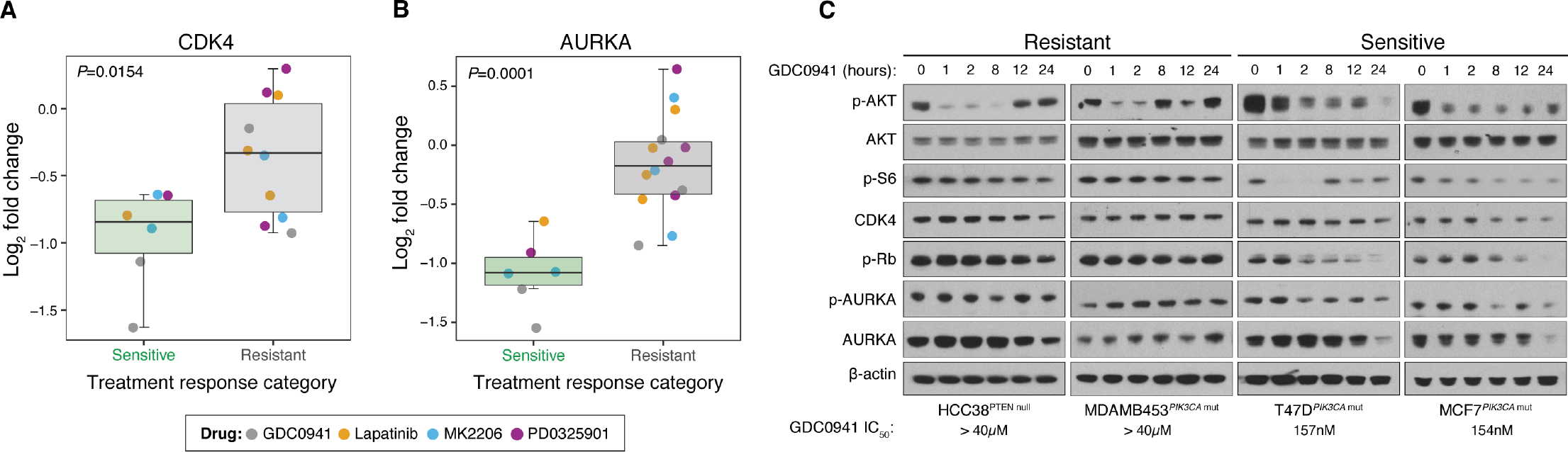
Maintenance of CDK4 and AURKA levels is associated with resistance to PI3K inhibition. (**a**) Changes in activity of CDK4 as measured by MIBs in drug sensitive versus drug resistant treatment responses after 24 hours of exposure to the indicated compounds. Each point reflects a single cell line and drug treatment. (**b**) Changes in activity of AURKA in drug sensitive versus drug resistant treatment conditions. (**c**) Western blot showing PI3K, CDK4, and AURKA signaling in GDC-0941-resistant and GDC-0941-sensitive cell lines. Protein lysates from cells treated with 1μM GDC-0941 were extracted at different points time, separated by SDS-PAGE, and analyzed by immunoblot with the indicated antibodies. Whiskers span the 25-75 percentiles. *P* values calculated using a two-sided t-test.

Analysis of dynamic responses across the MIBs screening panel revealed a significant association between maintenance of AURKA activity and drug resistance as the top hit (Fig. 2b). AURKA regulates centrosome alignment, mitotic spindle formation and chromosome segregation during mitosis^31^. Its activity and abundance are regulated by a wide range of factors involved in mitotic progression and microtubule assembly, and contributes to survival in the presence of mitotic abnormalities. In response to treatment with GDC-0941 in the panel of sensitive and resistant breast cancer cell lines, we observed a decrease in the abundance of total AURKA as well as its activity based on monitoring AURKA auto-phosphorylation in sensitive cells, whereas resistant cells maintained their AURKA levels and activity throughout the course of treatment (Fig. 2c, Supplementary Fig. 5f,g). The association of AURKA suppression with drug sensitivity also held true for the AKT inhibitor, as we observed a decrease in AURKA and phospho-AURKA levels only in MK2206 sensitive cells (Supplementary Fig. 5h-j). These results confirm that failure to suppress AURKA activity is associated with resistance to the PI3K inhibitor GDC-0941 and the AKT inhibitor MK2206 in breast cancer cells.

### CDK4 inhibition enhances sensitivity to PI3K-pathway inhibitors

We next asked if the identified candidates limit the efficacy of PI3K-pathway directed therapy. Since maintenance of CDK4 activity was associated with drug resistance, we tested whether CDK4 inhibition was sufficient to confer sensitivity to PI3K-pathway inhibitors. To assess this, we used a drug combination profiling approach to measure synergistic effects on cell viability across an extended panel of 13 breast cancer cell lines using CDK4 inhibitors and PI3K-pathway targeted therapies. To test drug combinations, we applied a dose matrix of increasing concentrations of the CDK4/6 inhibitor LEE011 alone and in combination with either a PI3K (GDC-0941), AKT (MK2206), or mTOR (RAD001) inhibitor and measured effects on cell proliferation. To evaluate drug synergy we used three methods: (1) visualization of Loewe excess which measures the increase in drug sensitivity over an additive model of the two drugs^32, 33^, (2) the combination index which measures shifts in drug potency with CI values < 1 indicating synergistic combinations^34^, and (3) the synergy score calculation which is a weighted sum of Loewe excess values^35^. Our results in MCF7 cells indicated that LEE011 in combination with GDC-0941, MK2206 or RAD001 is synergistic based on analysis of cell proliferation using Loewe excess, combination index and drug synergy score (Fig. 3a, Supplementary Fig. 6). Testing the combination with GDC-0941 across the extended panel of cell lines we found significant synergy in 62% of models (8 out of 13) based on a synergy score > 1, which we determined through simulation to represent a less than 5% chance of non-synergy (i.e. FDR < 5%) (Fig. 3b, Supplementary Fig. 6, see Methods). These findings were significant in light of previous observations of synergy between CDK4/6 and PI3K inhibitors^20, 36^, although it is unclear the extent to which this applies using other inhibitors of the PI3K-pathway. We therefore extended this analysis to drug combinations of LEE011 with either MK2206 or RAD001 and found significant synergy in 38% and 85% of models, respectively (Fig. 3b). We assessed whether synergy to these combinations might be selective to individual mechanisms of PI3K-pathway activation and/or breast cancer subtype. Overall we found no significant trend towards synergy based on *PIK3CA* or *PTEN* mutational status as well as receptor subtype in our combination studies, although this may be due to the composition of the cell line panel used in this study (Supplementary Table 2).

**Figure 3.**
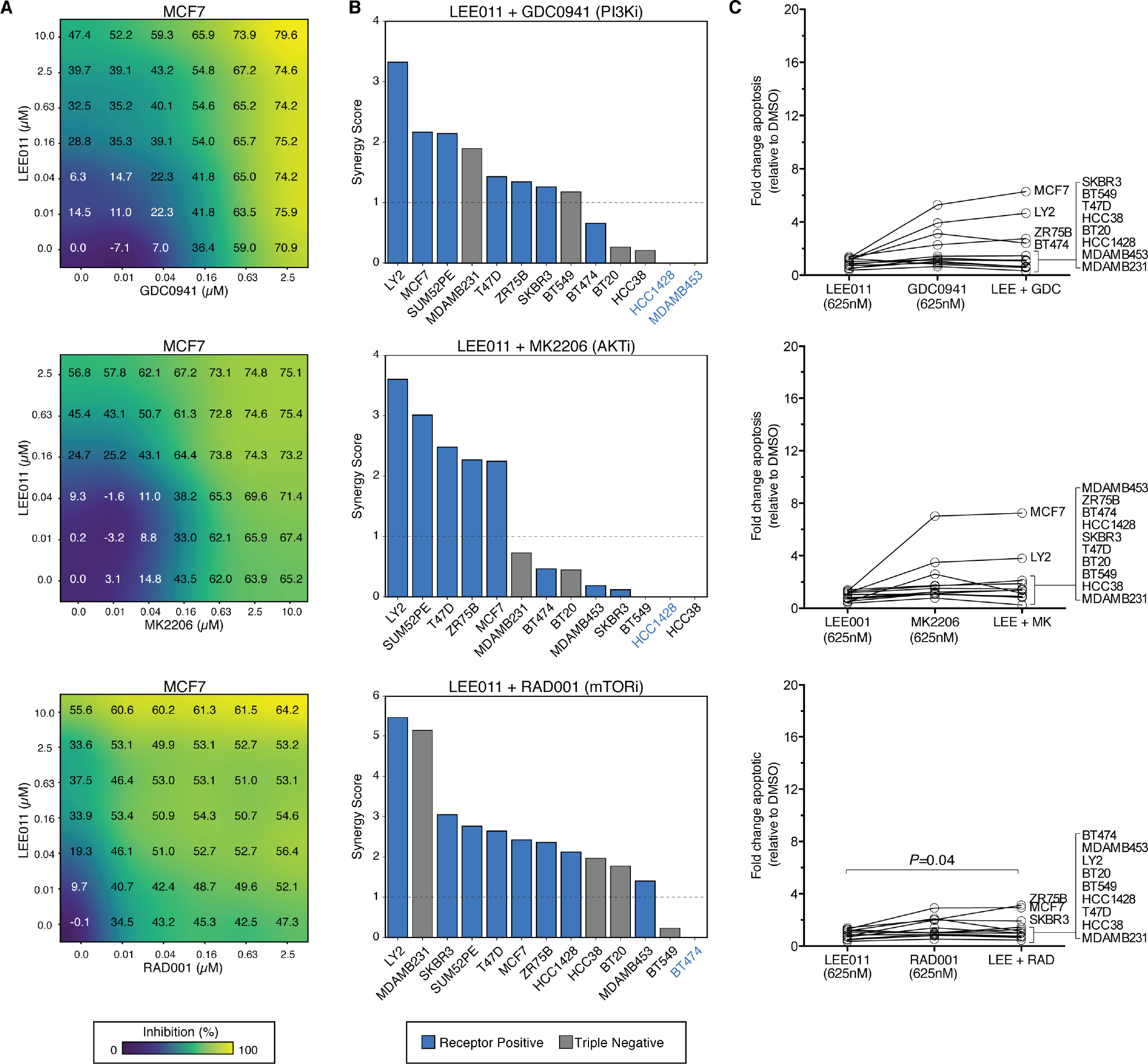
CDK4 suppression enhances sensitivity to PI3K-pathway inhibitors in breast cancer cell lines. (**a**) A dose matrix of GDC-0941 (PI3K), MK2205 (AKT), or RAD001 (mTOR) in combination with the CDK4/6 inhibitor LEE011 in MCF7 cells. Cell proliferation was assessed after 72 hours. Percent inhibition at each dose is shown. (**b**) 13 breast cancer cell lines were treated with the indicated combination in dose matrices across 6 concentrations for each agent and assessed for proliferation after 72 hours. The response was scored for synergy based on a Loewe excess inhibition model and ranked by synergy score. Dashed line at score of 1 indicates a 5% FDR cutoff to define synergistic combinations (see Methods). (**c**) Cell lines were treated with 625nM of the indicated single agents or combined together for 72 hours and apoptosis measured by YO-PRO1 positivity. Data are based on 4 replicates. Error bars are s.d. *P* values calculated using a two-sided t-test.

Since PI3K-pathway inhibitors are primarily cytostatic^2^, we next sought to determine if the addition of CDK4 inhibitors was capable of inducing cytotoxic responses. We assessed cell death through measurement of YO-PRO-1 positivity, a marker of early apoptosis. Through combined treatment across 12 cell lines, we found that the addition of LEE011 did not cause an increase in apoptotic cell death (Fig. 3c), which was independent of the particular dose used (Supplementary Fig. 7a). Overall, we found that the addition of LEE011 to any PI3K-pathway inhibitor was often synergistic but failed to enhance apoptosis in nearly all cell lines (Supplementary Fig. 7b). PI3K-pathway inhibitors impinge on mTORC1 which regulates the activity of CDK4 through the translation of Cyclin D1 to drive cells into S-phase^19, 37, 38^. Consistent with a previous report^20^, our MIBs data and drug synergy analysis support a model whereby resistance to PI3K inhibitors is due to incomplete suppression of mTORC1 resulting in residual CDK4 activity, which can be fully blocked with the addition of CDK4/6 inhibitors such as LEE011. Beyond PI3K inhibitors, our results further indicate that CDK4 suppression is also sufficient to confer sensitivity to AKT and mTOR inhibitors in most breast cancer cell lines. However, we found that CDK4 is only necessary for proliferation rather than tumor cell survival in the presence of PI3K-pathway inhibitors.

### AURKA is a survival factor in response to PI3K-pathway inhibition

Since maintenance of AURKA was strongly linked with drug resistance in our dataset, we next asked if it was required for survival in response to PI3K-pathway inhibition. We used dose-response analyses to characterize combinations of the AURKA-specific inhibitor MLN8237 with various PI3K-pathway inhibitors and measured effects on cell proliferation. Our results in MCF7 cells indicated that combinations of AURKA and PI3K-pathway inhibitors GDC-0941, MK2206 or RAD001 were synergistic based on analysis of cell proliferation using Loewe excess, combination index and drug synergy scoring (Fig. 4a, Supplementary Fig. 8). Testing these combinations across 13 cell lines, we found strong synergy based on a synergy score greater than 1 in 38%, 54% and 85% of models with either GDC-0941, MK2206 or RAD001, respectively (Fig. 4b).

**Figure 4.**
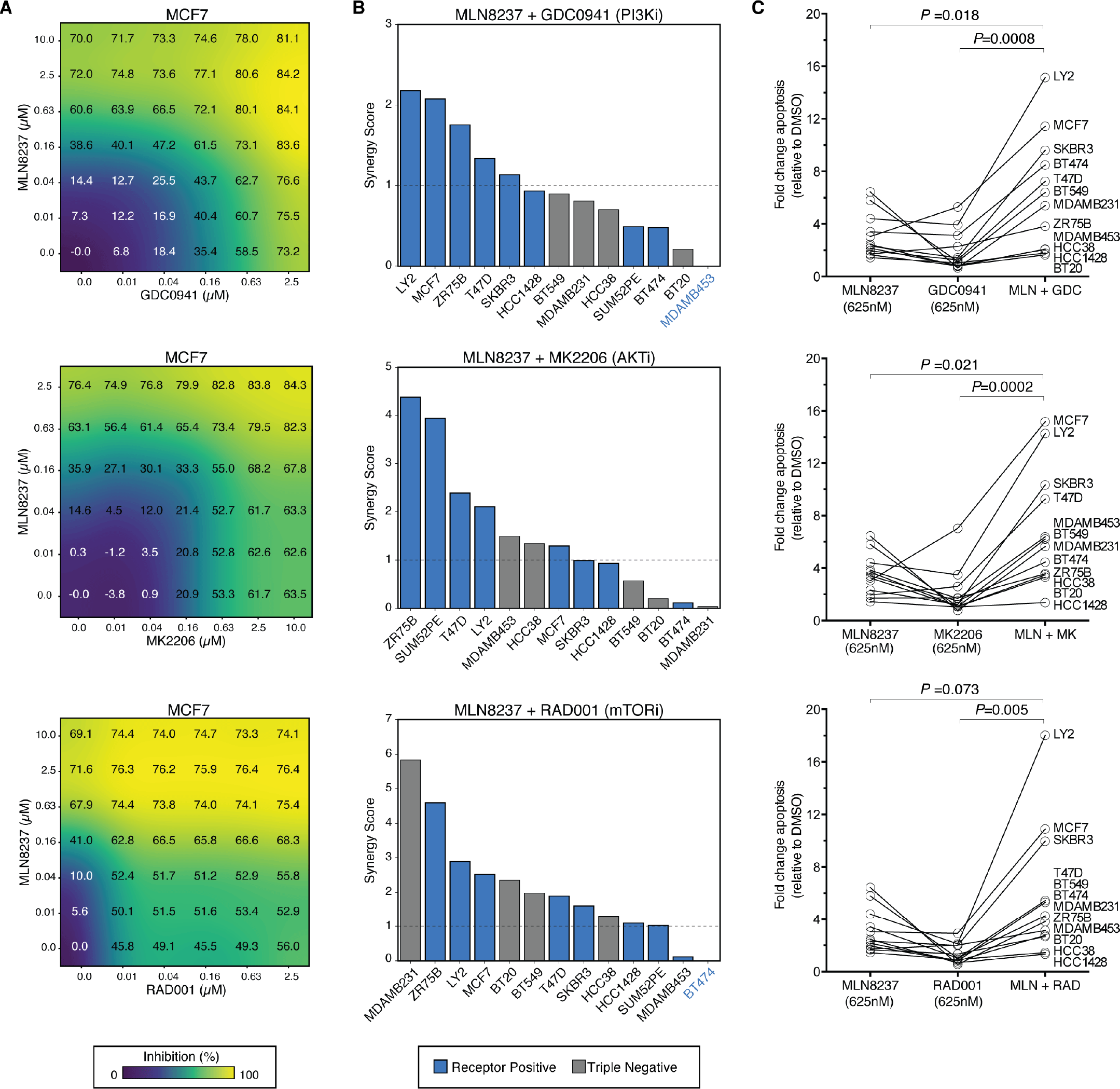
AURKA suppression enhances sensitivity and drives cell death in response to PI3K-pathway inhibitors in breast cancer cell lines. (**a**) A dose matrix of GDC-0941 (PI3K), MK2205 (AKT), or RAD001 (mTOR) in combination with the AURKA inhibitor MLN8237 in MCF7 cells. Cell proliferation was assessed after 72 hours. Percent inhibition at each dose of the drug is shown. (**b**) 13 breast cancer cell lines were treated with the indicated combination in dose matrices across 6 concentrations for each agent and assessed for proliferation after 72 hours. The response was scored for synergy based on a Loewe excess inhibition model and ranked by synergy score. Dashed line indicates a 5% FDR cutoff to define synergistic combinations (see Methods). (**c**) Cell lines were treated with 625nM of the indicated single agents or combined together for 72 hours and apoptosis measured by YO-PRO1 positivity. Data are based on 4 replicates. Error bars are s.d. *P* values calculated using a two-sided t-test.

Due to strong links between AURKA and the regulation of apoptosis^31^, we next asked whether AURKA inhibition could enhance cytostatic responses to PI3K-pathway inhibitors by converting them into cytotoxic effects. Notably, the addition of MLN8237 to GDC-0941, MK2206, or RAD001 resulted in an increase in apoptosis across cell lines, indicating that AURKA mediates cellular survival in the context of PI3K-pathway inhibition (Fig. 4c, Supplementary Fig. 7c). In many cases combinations including MLN8237 induced apoptosis greater than either single agent alone and this enhancement in apoptosis generally occurred in conditions where synergy was also observed (Supplementary Fig. 7d). In contrast to our findings with CDK4 inhibition, these data suggest that AURKA suppression is sufficient to sensitize cells to PI3K-pathway inhibitors through the induction of cell death. Therefore, AURKA is a promising companion target that may enhance the efficacy of PI3K-targeted inhibitors, warranting further validation of these findings in relevant preclinical models of breast cancer.

### Suppression of AURKA induces sensitivity to Everolimus (RAD001) by inducing cell death *in vivo*

Given the potent synergy and cytotoxicity engendered by co-targeting AURKA and the PI3K-pathway, we next evaluated the efficacy of this combination *in vivo.* For these studies we focused on the combination of MLN8237 with the only FDA-approved inhibitor targeting this pathway, the mTOR inhibitor RAD001 (Everolimus). Clinically, RAD001 overwhelmingly results in disease stabilization rather than remission^39, 40^. This is reflected *in vitro* where the response of MCF7 cells to RAD001 is cytostatic based on a high E_max_ of 0.54 and failure to induce PARP cleavage at high doses despite being highly sensitive (EC_50_ < 3nM) (Supplementary Fig. 9). To investigate whether AURKA suppression enhances response to RAD001 treatment in relevant preclinical systems, we generated orthotopic tumor models of breast cancer in immunodeficient mice. Consistent with a cytostatic effect, single-agent RAD001 treatment only partially impaired tumor growth over a period of 15 days. While single-agent MLN8237 also only partially blocked tumor progression, the combination of the two showed a significantly greater inhibition of tumor growth as compared to either single agent alone (*p* = 4.9e-9 compared to MLN8237, and *p* = 2.5e-5 compared to RAD001, Fig. 5a). Furthermore, all of the animals in the combination arm (n = 9/9) showed marked tumor regression, while no regressions were observed in either single agent arm (n = 0/13 in total, *p* = 2e-6 by Fisher’s exact test, Fig. 5b). Analysis of tumor specimens indicated an induction of apoptosis specific to the combination as demonstrated by an increase in the number of TUNEL-positive cells (Fig. 5c,d). During the course of study we did not observe any significant weight loss in the combination arm as compared to the RAD001 single-agent group (Supplementary Fig. 10), suggesting tolerability and no added toxicity from co-inhibiting Aurora kinase A. Therefore, the addition of MLN8237 to RAD001 treatment results in tumor regression and a strong cytotoxic response *in vivo.*

**Figure 5.**
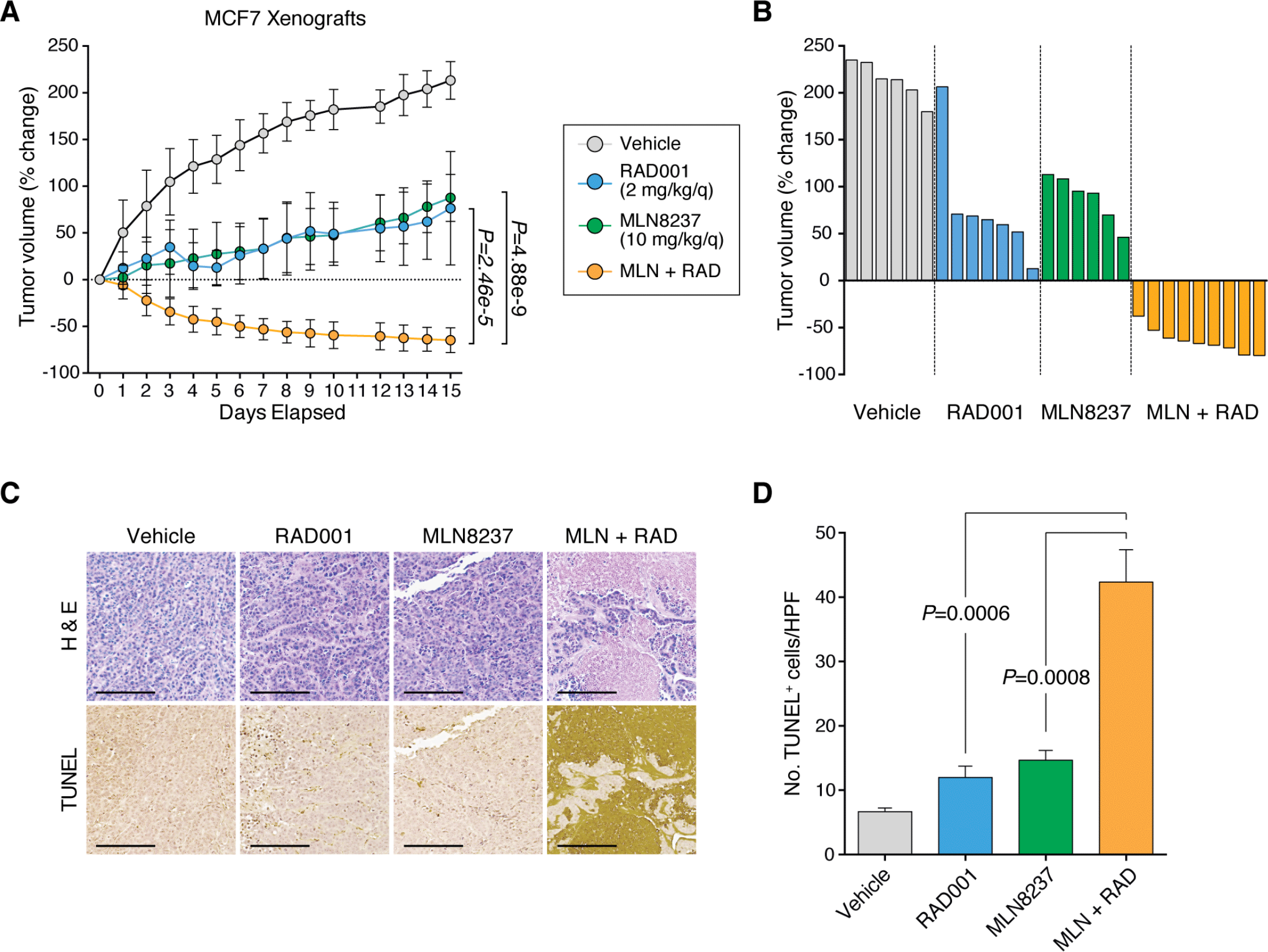
The Aurora kinase inhibitor MLN8237 enhances sensitivity to Everolimus (RAD001) and induces cell death *in vivo.* (**a**) MCF7 orthotopic xenograft tumors were treated with vehicle (n = 6 mice), RAD001 (2 mg/kg/day, n = 7 mice), MLN8237 (10 mg/kg/day, n = 6 mice) or the combination of the two single-agents (n = 9 mice) via oral gavage, daily, over 15 days. The percentage change in tumor volume was calculated for each animal from baseline. *P* values calculated using a two-sided t-test. (**b**) Individual tumor profiles compared to baseline for each tumor treated with vehicle, single agent RAD001, MLN8237, or the combination over a 15-day period. (**c**) Representative images of tumor tissue extracted from mice after 15 days treatment with the indicated agents and stained for H&E and TUNEL. Images shown using a 10x objective. Scale bars represent 200 *μ*m. (**d**) Quantification of the number of TUNEL^+^ cells/field (five high-powered (20x) fields from separate areas of each tumor) from TUNEL staining of MCF7 tumors following 15 days of treatment (n = 3 mice analyzed per treatment arm). *P* values calculated using a two-sided t-test. All error bars are s.d.

### Aurora kinase co-inhibition durably suppresses mTORC1 signaling

Because of observed synergy and tumor regressions *in vivo,* we next turned to identify mechanisms driving the increased efficacy of the drug combination. Since most PI3K-pathway inhibitors (including rapamycin derivatives such as RAD001) elicit feedback signals resulting in incomplete suppression of mTOR and consequent drug resistance^6,8, 19, 20^, we first asked if the combination of MLN8237 enhanced the activity of RAD001 on mTOR signaling to effectors involved in cap-dependent translation, S6 and 4E-BP1, *in vivo.* While we observed an incomplete and partial suppression of S6 in RAD001-treated tumors, the addition of MLN8237 resulted in a durable and complete loss of S6 in all 9 tumors (Fig. 6a). It is well established that while rapalogs such RAD001 are relatively potent inhibitors of the mTORC1 target S6, they are inefficient inhibitors of another mTORC1 target 4E-BP1 and therefore only partially impair cap-dependent protein synthesis^41^. To test whether the addition of MLN8237 might synergize with RAD001 by enhancing suppression of mTORC1, we investigated the activity of phospho-4E-BP1, a rapamycin-resistant output of mTOR that is stimulated by rapamycin treatment^42^. While phospho-4E-BP1 levels were enhanced with RAD001 single-agent treatment, co-treatment with MLN8237 suppressed these levels back to nearly baseline (Fig. 6a). This surprising finding led us to ask how Aurora kinase inhibition might alter this key signaling output of mTOR. For this purpose we investigated AKT activity via phosphorylation of serine 473, which activates mTOR and is catalyzed by a variety of kinases^43, 44^ (Fig. 6a). Single-agent MLN8237 reduced phospho-AKT levels both in monotherapy and combination treatment settings, indicating that Aurora kinases sustain mTOR levels by promoting AKT activity. We next examined whether this Aurora kinase driven maintenance of mTOR was a feature of RAD001 or a general feature of PI3K-pathway inhibitors. Using MCF7 cells *in vitro,* we observed that single-agent MLN8237 was sufficient to impair phospho-AKT and that the combination of MLN8237 with either GDC-0941 (targeting PI3K) or MK2206 (targeting AKT) led to robust ablation of phospho-S6 and phospho-4E-BP1 levels (Fig 6b). Therefore, Aurora kinases contribute to resistance to PI3K-pathway inhibitors through the maintenance of AKT activity, which drives residual mTORC1 dependent protein synthesis. Targeting this Aurora kinase dependent survival mechanism using the combination of MLN8237 and PI3K-pathway inhibitors results in a more durable and complete repression of mTORC1 activity.

**Fig. 6.**
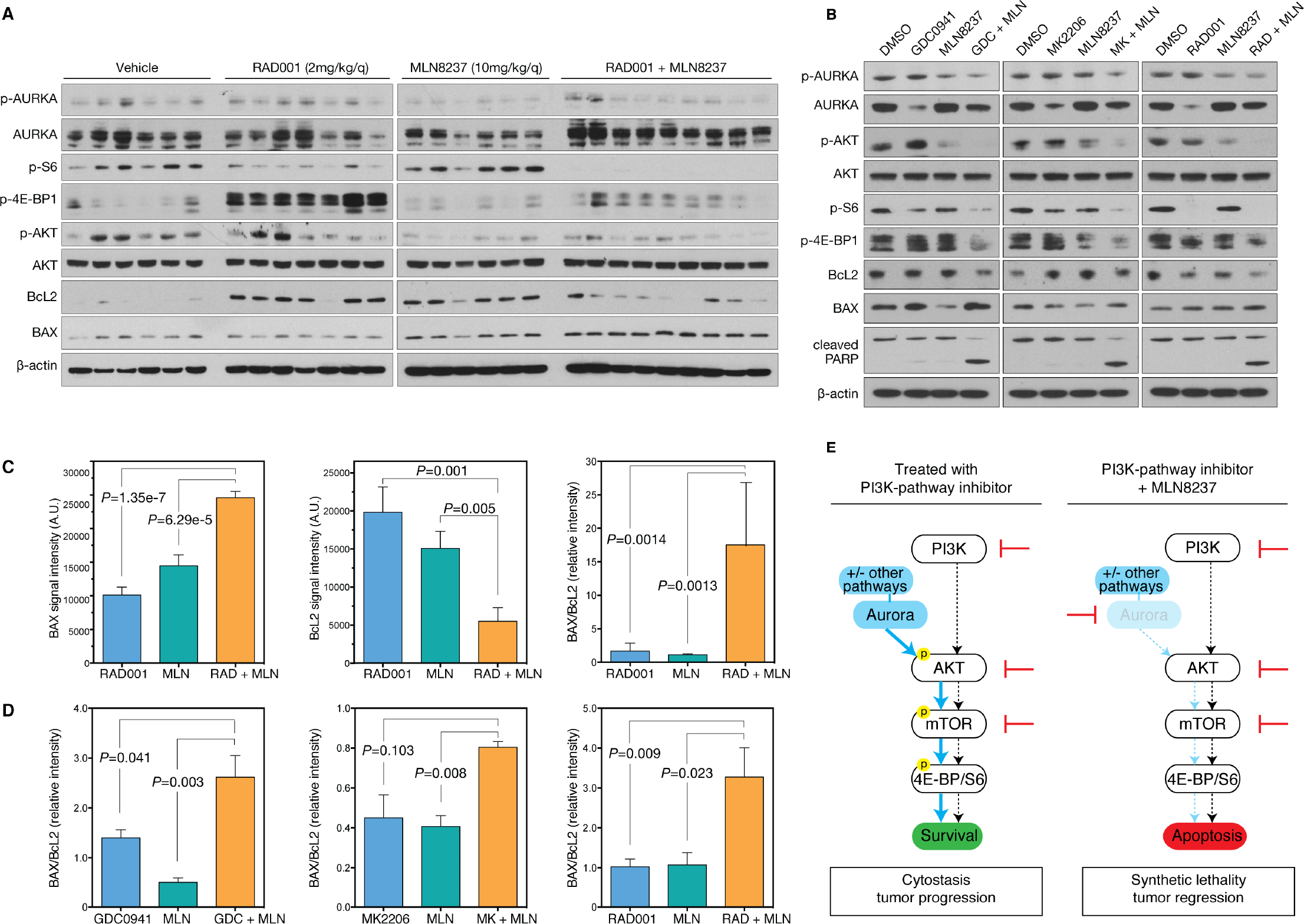
Aurora kinase co-inhibition durably suppresses mTORCI signaling and alters the Bax/BcL2 ratio. (**a**) MCF7 orthotopic xenografts were treated with vehicle (n = 6 mice), RAD001 (2 mg/kg/day, n = 7 mice), MLN8237 (10 mg/kg/day, n = 6 mice) or the combination of the two single-agents (n = 9 mice) for 15 days, at which point tumors were harvested and snap frozen. Western blot of protein lysates from individual tumors were probed with the indicated antibodies. (**b**) MCF7 cells were treated with either 250nM GDC-0941, 250nM MK2206, 5nM RAD001, 100nM MLN8237 or the indicated combinations for 24 hours and protein lysates subjected to immunoblot using the indicated antibodies. (**c**) BAX, BcL2 and BAX/BcL2 ratio in MCF7 orthotopic xenografts treated for 15 days with the indicated drugs based on quantification of western blot images. (**d**) BAX/BcL2 ratio in MCF7 cells treated for 24 hours with the indicated drugs based on quantification of western blot images. Error bars based on a minimum of 3 replicate experiments. (**e**) Proposed model of mechanism of Aurora kinase inhibitor synergy. *De novo* resistance to single agent inhibition of PI3K, AKT or mTOR is due to incomplete suppression of the pathway due to Aurora kinase signaling which activates AKT. Drug combinations that simultaneously inhibit the PI3K-pathway and block Aurora kinase signaling completely suppress mTOR signaling to 4E-BP1 and S6 resulting in tumor cell death and synthetic lethality. Error bars are s.e.m. *P* values were calculated using a two-sided t-test.

### Aurora kinase co-inhibition leads to apoptosis through unbalancing pro-and anti-apoptotic factors

Cap-dependent protein synthesis is critical for cell growth and survival, and drug combinations led to simultaneous inhibition of 4E-BP1 and S6 and were associated with cell death, as evidenced via PARP cleavage *in vitro* (Fig. 6b), and TUNEL positivity *in vivo* (Fig. 5d). Therefore, we next sought to characterize the mechanism by which Aurora kinase mediates cell survival in response to PI3K-pathway suppression. Since both Aurora kinases and mTOR regulate a number of components of the intrinsic apoptosis pathway^31, 45^, we hypothesized that deregulation of the balance of pro-and anti-apoptotic factors may cause cell death in response to drug combinations containing MLN8237. BcL2 family-member BAX promotes apoptosis through permeabilization of the mitochondrial outer membrane, while BcL2 prevents apoptosis by inhibiting the activity of BAX. Together, the balance of these two proteins forms a molecular rheostat for apoptosis^46^. In xenografted tumors, combination treatment resulted in increased BAX levels and a reduction in BcL2 levels leading to an increase in the ratio of BAX/BcL2 compared to either MLN8237 or RAD001 treatment alone (Fig. 6c). We confirmed that this specific deregulation of apoptotic family members was a general feature of the pathway since the BAX/BcL2 ratio was also increased by the addition of MLN8237 to GDC-0941, MK2206 or RAD001 in MCF7 cells *in vitro* (Fig. 6d). Taken together, we propose a model whereby Aurora kinase inhibitors potentiate the activity of PI3K-pathway inhibitors through enabling a durable and complete suppression of AKT/mTOR signaling, and drive cell death by altering the balance of pro and anti-apoptotic factors (Fig. 6e).

## DISCUSSION

Through an unbiased proteomics approach to assay kinase activity, we measured dynamic changes elicited by therapy as a means to develop novel drug combinations. In contrast to previous work limited to a single drug and cell type^9, 22, 26^, the systematic measurement of kinome dynamics across a diverse set of cell lines allowed us to map molecular changes associated with resistance to a variety of inhibitors targeting core pathways in breast cancer. The largest and most unbiased dataset of its type, this data is a resource for the interrogation of signaling pathways modulated by drug treatment in breast cancer. Using this approach, we found a number of cases where failure to inhibit a particular kinase was associated with drug resistance. We hypothesize that many of these kinases may represent survival factors that limit the efficacy of therapy by modulating cellular dependence on the target pathway. Since our proteomic screen included multiple drugs that impinge on distinct oncogenic pathways, we found it surprising that a set of common survival factors were identified. This is likely due to the convergence of both the PI3K and MAPK pathways on a set of shared effectors including protein synthesis^17, 19, 47, 48^. For two of the factors amenable to pharmacologic inhibition, CDK4 and AURKA, we show that their presence limits the efficacy of PI3K-pathway targeted therapy and thus represent synthetic lethal targets to enhance therapeutic responses. Future work may determine if other candidates we identified also act as survival factors and how they might do so.

We hypothesized that mapping kinome dynamics would reveal survival factors whose maintenance contributed to drug resistance. In the case of CDK4 and AURKA, their suppression also appears to be sufficient to induce sensitivity to PI3K-pathway inhibitors. Our data are consistent with prior work showing that CDK4/6 inhibitors sensitize cells to both PI3K and HER2 inhibitors, findings which have precipitated multiple clinical trials (NCT02154776, NCT02088684, NCT01872260, NCT02389842)^20, 36, 49,50,51–52^. Here we extend these findings to other inhibitors of the pathway to include those targeting AKT and mTOR, suggesting that CDK4 may act as a molecular barrier to the efficacy of a wide variety of drugs that impinge on the activity of the PI3K-pathway. However, combinations including CDK4/6 inhibitors only impaired cell proliferation rather than survival. As clinical trials testing CDK4/6 inhibitor combinations are ongoing, it remains to be seen the impact this distinction will play on patient responses.

We determined that maintenance of AURKA was strongly associated with drug resistance, suggesting that AURKA suppression is necessary for drug sensitivity. Maintenance of AURKA was also sufficient to confer drug resistance in a variety of cell lines as evident by the widespread drug synergy we observed. We show that in response to treatment with PI3K-pathway inhibitors, Aurora kinases maintain the activation of AKT and drive residual mTOR activity. Co-inhibition of Aurora kinases with MLN8237 fully blocks this residual mTOR activity, impairing protein synthesis and resulting in cell death. These findings also highlight the importance of AKT activation through serine 473 as a route of drug resistance. While a number of kinases have been shown to operate at this site including PDK2, ILK and mTOR though feedback via mTORC2^52^, it remains unclear whether Aurora kinases act on this site directly or indirectly through these factors. Future work could also investigate how AURKA activity is maintained in resistant settings, although this will likely be complex since AURKA is regulated by multiple interacting factors that govern its abundance and activation during mitosis and interphase^53^. Although this aspect of its biology is poorly understood, the PI3K-pathway has been reported to influence mitotic progression^54, 55^, and future work could explore how it might participate in the regulation of AURKA. Therefore, AURKA acts as a molecular barrier to the efficacy of PI3K-pathway inhibitors and mediates survival in response to treatment by maintenance of mTOR signaling.

Our findings reveal that the combination of Aurora kinase inhibitors and PI3K-pathway inhibitors is synergistic and could be a promising clinical strategy to enhance treatment response in breast cancer. Clinical data of PI3K and mTOR inhibitors have shown only modest benefit in breast cancers, with around 2-4 months improvement in PFS and largely disease stabilization in patients^39, 56^. Consistent with these clinical observations, most inhibitors in this class cause only a proliferative arrest *in vitro^57^* and it has been proposed that combinations that induce apoptosis may be used to enhance responses^58^. We observed strong synergy in most cell lines using MLN8237 and any of the PI3K-pathway inhibitors, particularly in addition to RAD001 treatment where we identified synergy in 85% of cell lines. In contrast to cytostatic combinations with the CDK4/6 inhibitor (i.e. synthetic sickness), we found that combinations with Aurora kinase inhibitors were synthetic lethal and potently induced cell death. As single-agent therapies, Aurora kinase inhibitors have reached phase 3 clinical trials for lymphoma with manageable toxicities, albeit limited efficacy when used as monotherapies^59^. Given that the most common adverse events of PI3K-pathway inhibition are hyperglycemia, rash, and gastrointestinal toxicity, and those of Aurora kinase inhibition are primarily neutropenia, we are encouraged that the non-overlapping toxicity profile between the two agents may be well tolerated in patients as they were in our *in vivo* studies. As single-agent responses to both PI3K-pathway and Aurora kinase inhibitors have been modest, these findings may unlock the potential of these agents in realizing a clinical benefit. Therefore, a further study of these drug combinations in the clinical setting is warranted.

## ONLINE METHODS

### Breast cancer cell lines and reagents

BT549, BT20, and SKBR3 cells were obtained from the UCSF Cell Culture Facility. BT474, HCC1428, HCC38, LY2, MCF7, MDAMB231, MDAMB453, T47D, SUM52PE, and ZR75B cell lines were obtained from the American Type Culture Collection (ATCC). Lines were grown according to published protocols^60^ except for SKBR3 which was cultured using RPMI media supplemented with 10% fetal bovine serum (FBS) and 1% pen/strep. All cell lines tested negative for mycoplasma contamination. Drugs used in this study were purchased from Selleck Chemicals (GDC-0941, MK2206, PD0325901, Lapatinib, MLN8237, and LEE011) and LC Laboratories (RAD001).

### MIBs analysis

Multiplexed inhibitor bead enrichment and MS analysis (MIB/MS) were performed as described previously^9, 22, 24^. In summary, a selection of bait compounds were purchased or synthesized and coupled to sepharose using 1-Ethyl-3-(3-dimethylaminopropyl)carbodiimide–catalyzed chemistry. After 24-hour treatment with drug or DMSO, cell lysates were diluted in binding buffer with 1 mol/L NaCl and kinase enrichment was performed using gravity chromatography following pre-clearing. After washing, the bound kinases were eluted with SDS followed by extraction/precipitation, tryptic digest and desalting. Liquid chromatography-tandem mass spectrometry (LC/MS-MS) was performed on a Velos Orbitrap (Thermo Scientific) with in-line high-performance liquid chromatography (HPLC) using an EASY-spray column (Thermo Scientific). Peptide identifications were made using ProteinProspector (v5.10.10) and input into Skyline for label-free quantification^61^.

Peptide quantification data were pre-processed before analysis with MSstats v2.3.3^62^. First, library peptides and peptides that map to non-kinase proteins were removed. Kinase peptide peak area values were log_2_-transformed and quantile-normalized to correct for variation between replicates. Finally, peptides that mapped to multiple kinases were removed, as well as peptides that were entirely missing in one or more conditions. For each kinase, the log_2_ ratio of each drug-treated condition to the DMSO control was estimated using the mixed-effects regression model in MSstats.

### Drug combination studies

Cell lines were seeded in 384-well assay plates at a density of 1,000 cells/well in a total volume of 40 *μ*L/well, and incubated at 37°C, 5% CO_2_ overnight. Dose matrices were assembled containing 6-point, 4-fold serial dilutions from the top concentration for each agent on the x-and y-axes. Following 72 hours of drug exposure, proliferation and cell death was measured by staining with Hoescht (Life Technologies) nuclear dye and YO-PRO-1 (Life Technologies), respectively, and analyzed using a Thermo CellInsight High Content microscope. Raw phenotype measurements from each treated well were normalized to the median of vehicle-treated control wells and examined for synergistic effects between both compounds.

To evaluate drug combinations we used a Loewe model of drug additivity and calculated a synergy score. First, we fit a sigmoidal function to each of the single agent responses. Next we calculated the expected inhibition for each combination using the Loewe additivity model^32^. The synergy score *s* was calculated as previously defined^35^ as a positive-gated inhibition-weighted volume over of Loewe additivity:

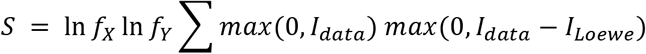

Where *f_x_* and *f_Y_* are the dilution factors used for compounds *X* and *Y* respectively, *l_data_* is the matrix of inhibition data at this dilution factor, and *I_Loewe_* is the expected inhibition according to Loewe additivity. *CI_50_* values for equal-dose combinations were calculated as previously defined^34^:

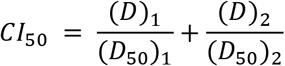

Where (*D*)_1_ and (*D*)_2_ are the given doses of the two drugs, and (*D*_50_)_1_ and (*D*_50_)_2_ are the *IC*_50_ values for each drug as a single agent.

To determine a cutoff for the synergy score we simulated the distribution of scores generated by an additive drug combination. We generated two hypothetical compounds by sampling random shape parameters for their dose-response functions, and calculated the expected Loewe model of the combination. We then added normally distributed noise to the model with variance estimated from our experimental data and calculated the resulting synergy score. This process was repeated 100,000 times to simulate the distribution of synergy scores for different additive combinations. The 95^th^ percentile of this distribution was 0.91 and so we conservatively identified combinations with *S* ≥ 1 as synergistic.

### Western blotting and antibodies

Proteins were extracted using RIPA buffer (50 mM Tris-HCl pH 7.5, 150 mM NaCl, 0.1% sodium deoxycholate, 0.1% SDS, 1 mM EDTA pH 8.0, 1% NP-40) containing proteinase (Roche) and phosphatase (Roche) inhibitor cocktails. Samples were resolved using 4-12% SDS-PAGE gels (Life Technologies) and transferred to PVDF membranes (Millipore). Membranes were probed overnight on a 4°C shaker with primary antibodies (1:1,000 dilution unless indicated) recognizing the following proteins: p-AKT (Ser473) (9271, Cell Signaling), AKT (4691, Cell Signaling), p-S6 (Ser240/244) (5364, Cell Signaling, 1:20,000), p-4E-BP1 (Thr37/46) (2855, Cell Signaling), CDK4 (12790, Cell Signaling), p-Rb (Ser780) (9307, Cell Signaling), p-AURKA (Thr288) (3079, Cell Signaling), AURKA (4718, Cell Signaling), Cleaved PARP (Asp214) (9541, Cell Signaling), BcL2 (2870, Cell Signaling), BAX (2772, Cell Signaling), and β-actin (3700, Cell Signaling).

### Mouse xenograft studies

All animal studies were conducted in accordance with the UCSF Institutional Animal Care and Use Committee (IACUC). 4-week old immunocompromised NOD/SCID female mice were purchased from Taconic Biosciences, and MCF7 cells used for *in vivo* transplant were obtained from the UCSF Preclinical Therapeutics Core. Xenograft tumors were initiated in the cleared mammary fat pads of mice bearing slow release estrogen pellets (Innovative Research of America) by orthotopic injection of 1e6 MCF7 cells in a 1:1 mixture of serum-free medium and Matrigel (BD Biosciences). When tumors reached ≥ 1 cm in any direction via electronic caliper measurements, mice were randomized into cohort groups and treatment was initiated.

Treatment arms received either vehicle (1:1 mixture of single-agent diluents), RAD001 formulated as a microemulsion (2mg/kg/q; 30% Propylene glycol, 5% Tween 80), MLN8237 (10mg/kg/q; 10% 2-hydroxypropyl-β-cyclodextrin, 1% sodium bicarbonate), or the combination daily, via oral gavage. Animals were monitored daily for evidence of toxicity including weight and skin effects, and changes in tumor size (mm^3^) through bidirectional measurements of perpendicular diameters using electronic calipers, and calculated as *V* = 1/2(*length* × *width*^2^). Mice were sacrificed after 15 days of treatment, following which tumors were excised and a portion of the tissue fixed in 4% paraformaldehyde. The remaining tumor tissue was flash-frozen in liquid nitrogen.

### Immunohistochemical analysis

PFA-fixed tumor samples were paraffin-embedded, and immunohistochemical staining of tissue sections was performed. TUNEL staining was carried out using the ApopTag Peroxidase *In situ* Apoptosis Detection Kit (Millipore), according to the manufacturer’s instructions (n = 3 tumors per experimental group). Stained slides were digitized using the Leica DMi1 Microscope (Leica Microsystems) with a 20x objective. Images were scored as the number of TUNEL-positive cells per captured field, and quantification was performed in a manner that was blinded to treatment group.

### Statistical analysis

Data are expressed as means ± s.d., unless otherwise indicated. Statistical analyses were performed using GraphPad Prism 6 (version 6.0g) and R (version 3.32). Two-tailed Student *t* tests (with unequal variance) were used in all comparisons unless otherwise noted. *P* < 0.05 was considered statistically significant throughout the study.

## Acknowledgments

The authors would like to thank members of the Bandyopadhyay laboratory for helpful discussions and technical assistance. We also thank Andrew Beardsley, Evan Markegard, Davide Ruggero and William Weiss for helpful discussions and reagents.

## Funding

This work was supported by NCI U01CA168370 and NIGMS R01GM107671 (S.B.), Martha and Bruce Atwater (S.B., A.G.), OHSU Pilot Project Funding (S.B., J.K.), and an American Cancer Society Postdoctoral Fellowship (J.G.).

## Author Contributions

Conceptualization: H.J.D., J.T.W., J.G., J.K., S.B.; Data analysis: H.J.D., J.T.W., J.G.; investigation: H.J.D., J.T.W., N.B., R.L., R.C., O.M., K.S.; writing of original draft: H.J.D., S.B.; manuscript finalization: all authors; funding: S.B., A.G.; supervision: S.B., A.G.

## SUPPLEMENTARY MATERIALS

Supplementary Figure 1. Structure of drugs bound to multiplex inhibitor bead column.

Supplementary Figure 2. Dependence of MIBs data on gene expression.

Supplementary Figure 3. Diversity analysis of breast cancer cell lines.

Supplementary Figure 4. Cell line dose response analysis and correlation with sensitivity.

Supplementary Figure 5. GDC-0941 and MK2206 validation in sensitive and resistant cell lines.

Supplementary Figure 6. Drug synergy analysis for combinations including the CDK4 inhibitor LEE011.

Supplementary Figure 7. Induction of apoptosis in selected single agent and drug combinations.

Supplementary Figure 8. Drug synergy analysis for combinations including the Aurora kinase inhibitor MLN8237.

Supplementary Figure 9. RAD001 is cytostatic in MCF7 cells.

Supplementary Figure 10. Mouse weight during xenograft studies.

Supplementary Table 1. Scores and p-values from MIBs profiling.

Supplementary Table 2. Synergy scores for combinations in this study.

